# Decoding Images from Multi-Region, High Resolution, Electrode Recordings In the Mouse Visual System

**DOI:** 10.1101/2021.07.07.451504

**Authors:** Chris Fritz

## Abstract

We hypothesize that deep networks are superior to linear decoders at recovering visual stimuli from neural activity. Using high-resolution, multielectrode Neuropixels recordings, we verify this is the case for a simple feed-forward deep neural network having just 7 layers. These results suggest that these feed-forward neural networks and perhaps more complex deep architectures will give superior performance in a visual brain-machine interface.

## 2 Introduction

A visual brain-machine interface needs to decode images from cortical activity [1], particularly natural images which have distinct spatial frequency spectra [2]. While natural images cause sparse activation of neurons in visual cortex (V1) [3] [4] [5], how this sparse set of neurons represents a given image and how the image can be inferred (decoded) from sparse activity is unclear.

Linear decoding assumes that the represented image is a linear function of neuron activity. Such decoders underlie classic visual neuroscience theory [6] [7], but are results of experiments featuring small numbers of electrodes, e.g. [8]. As advancements such as optogenetics, two-photon imaging, and electrode design increase the scale of availale physiological data, linear decoders may struggle to keep pace in their performance, while deep learning continues to outperform linear decoders as the state of the art in a multitude of tasks and brain regions [9] [10] [11] [12] [13]. The caveat to increased performance is that deep networks perform best with larger training data in the form of function input/output.

Until recently, direct neural data in the form of electrode recordings were limited to few (order of one hundred) measurements in a single brain region using hardware such as the Utah Array. [14] Neuropixels [15] are a recent hardware advancement that give not only an order of magnitude increase in number of electrodes, but also data from multiple brain regions simultaneously. A public dataset of neuropixel recordings in mice was recently made available online [16]. With this rapid expansion of available data, deep networks are poised to offer state of the art decoding performance from visual cortex activity.

CS230: Deep Learning, Winter 2018, Stanford University, CA. (LateX template borrowed from NIPS 2017.)

## 3 Data Collection & Feature Extraction

The Neuropixels Dataset [17] covers 58 experiments in which visual stimuli ranging from Gabor functions to natural scenes are presented to mice with multiple neuropixels probes inserted into visual cortex. The length of the probes also permits recordings from subcortical structures such as the Lateral Genticulate Nucleus (LGN) and Lateral Posterior (LP) nucleus. High-pass-filtered electrode recording data is statistically whitened before being passed through the Kilosort2 algorithm to identify specific neurons, giving spike times for a set of neurons/units assigned to a given anatomical region [18].

The decoder maps measured spiking rates recorded by the probes to the images presented to the subject on a 1920 × 1080 monitor. That is, we infer the image on the monitor from the spiking activity.

### Input Spike Rates

We compute the spike rates for each of *N*_*neurons*_ by measuring the number of spikes in window of length *τ* after a given image is presented as in figure (1).

**Figure 1:**
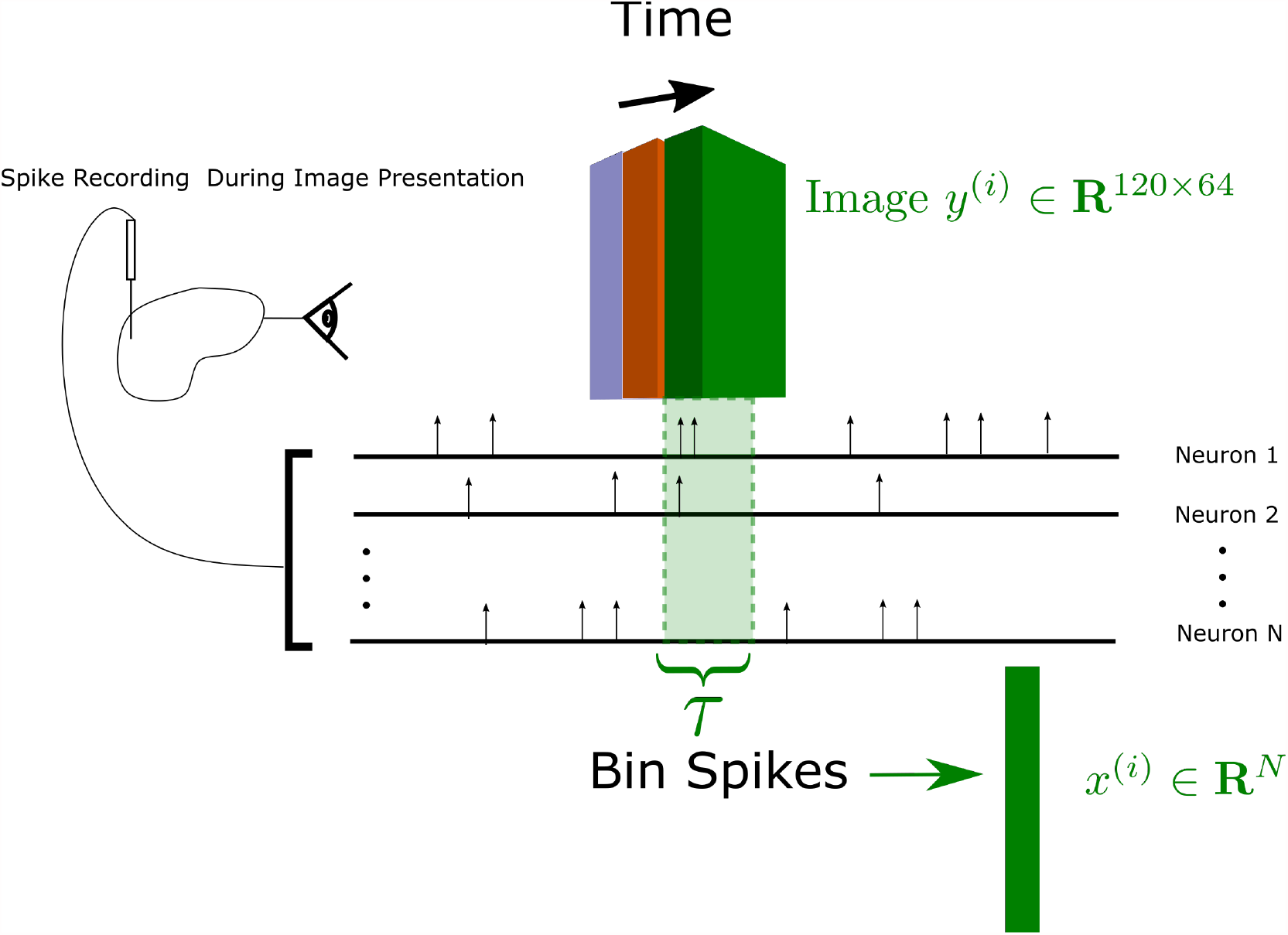
Computing Decoder Input: Spike rates are computed by counting the number of spikes in a window of length *τ* after the image onset. The resulting spike rates are then z-scored to zero-mean and unit variance to form the decoder input data.

Spike rates are then z-scored so that each channel has zero mean and unit variance across the time of recording. Using the linear model described below, we empirically determine that the optimal window length is 167*ms*. As shown in figure (2), this length offers the best trade-off between accuracy and latency, since the improvement on accuracy versus integration time switches from exponential to linear.

**Figure 2:**
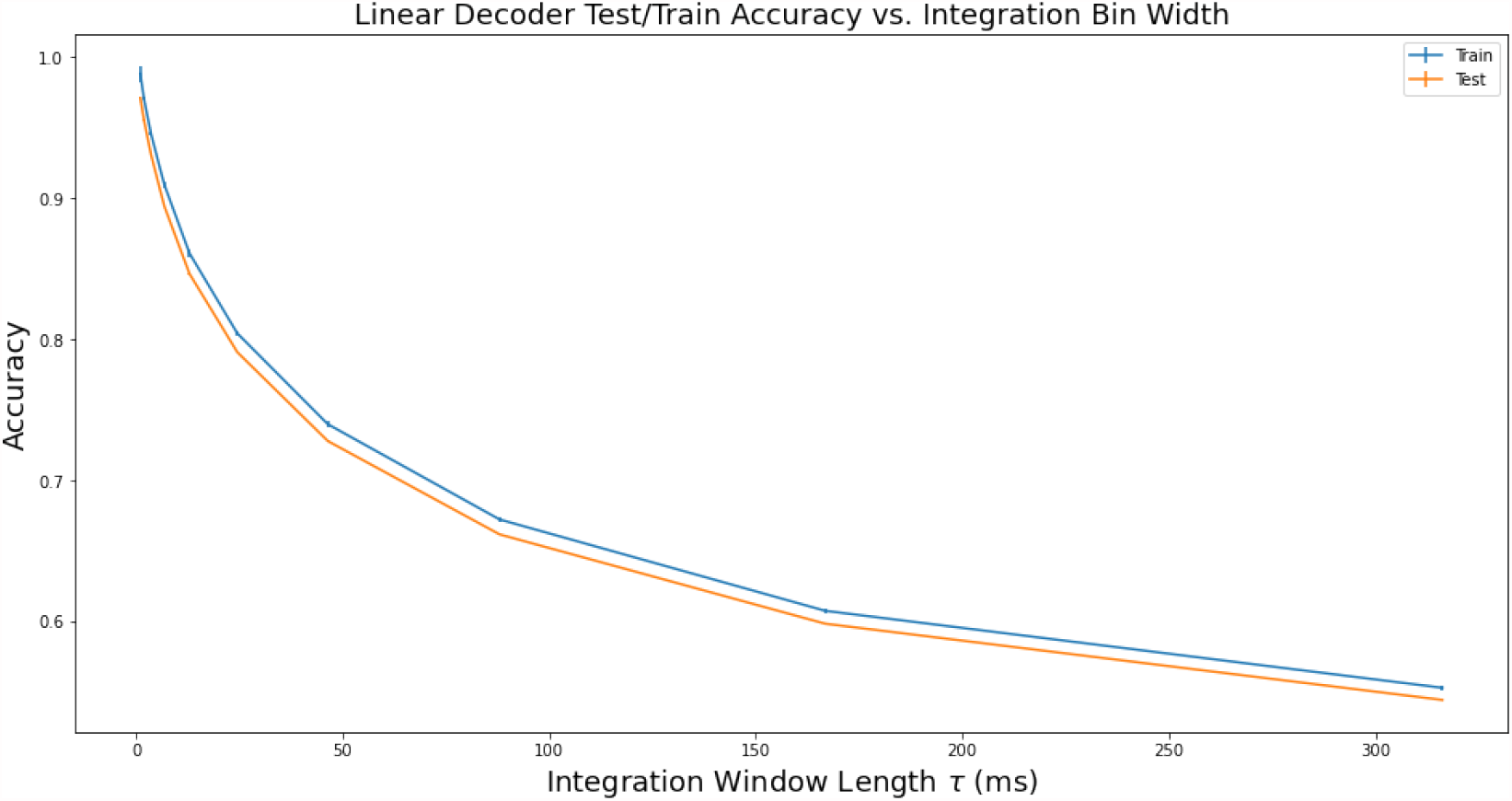
Determining the Optimal Integration Window length. Using a linear model, we computed the best least squares fit of spike rates to pixel intensities over 90% of the dataset. The remaining %10 percent of images was then predicted from the resulting linear model. The accuracy was computed using equation (2) over the resulting predictions.

### Output Images

The network’s output are the presented 1920 × 1080 images presented at the beginning of the integration window of length *τ* = 30 ms. To reduce the dimensionality, we resample the images by averaging to size 120 × 64. The monitor spanned 120° × 95° of the mouse visual field, so the resampled resolution limits the spatial frequency of the presented images to 0.5 *cycles/*°, the approximate limit of spatial frequency observable by mice [19]. Thus, we preserve all perceptible visual information while reducing the size of the image by a factor of 16^2^. To improve the network learning rate, we then zero mean the pixel intensities from [0, 255] to [−128, 127]. Finally, the images are flattened to 128 * 64 = 8192-vectors.

Here we considered recording data from one session (ID: 840012044) over exclusively natural stimuli (‘natural_movie_one_more_repeats’), giving a total of 54, 000 frames presented over 80 minutes at two separate intervals.

## 4 Decoding Model Selection, Training, & Evaluation

We evaluate a linear decoding model and compare it to a deep feed-forward neural network.

### Model Objective

We consider the mean square error as a loss function, common in image processing literature [20]. The mean square error (MSE) measures the norm of the difference in pixel intensities between the true image *y* and predicted image 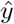:

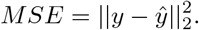

### Model Training

To train the deep network, we use batch gradient descent with the Adaptive Moment Estimation (Adam) algorithm [21] using a batch size of 1000, and a learning rate of *E* = .0001. The deep network model is a feedforward sequential network with 7 hidden layers with input length *N*_*pixels*_, depth of 7 and width of 77 as sketeched in figure (3). These numbers were chosen by training a new network for each combination of logarithmically spaced parameter values and evaluating the performance after 1000 epochs.

**Figure 3:**
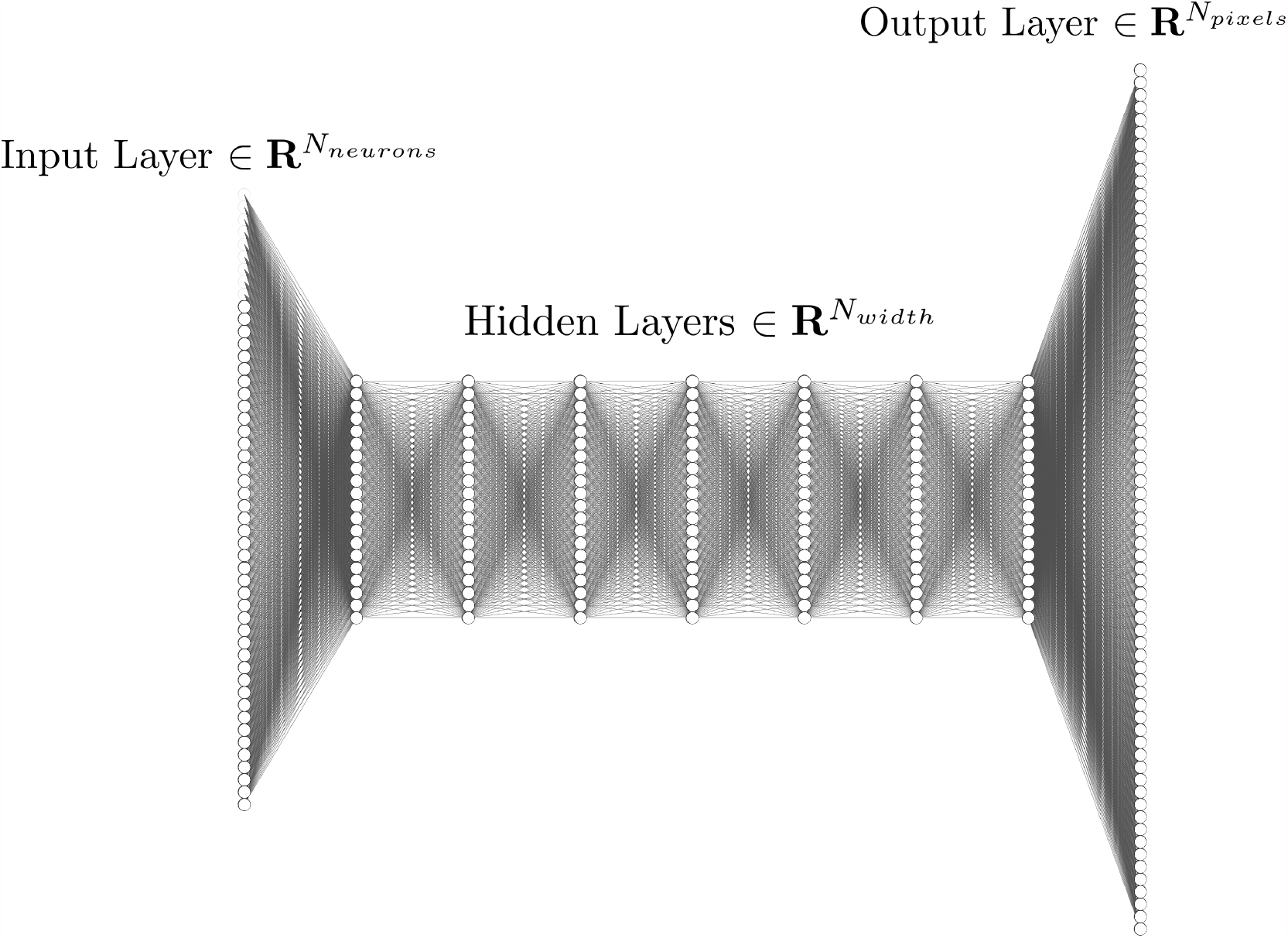
Sketch of the Feedforward Deep Network Used in Image Prediction. The parameters are *N*_*neurons*_ = 751, *N*_*width*_ = 77 with 7 layers, and *N*_*pixels*_ = 8192.

For the linear decoder, the MSE objective has an analytic solution:

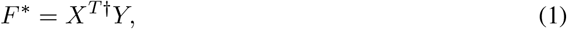

where 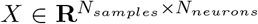, gives the recorded spike rates, 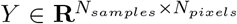 gives the presented images, and 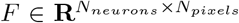, gives the linear mapping between spike rates and images. Each row of *F* gives the pixel intensities decoded by a spike in that respective neuron.

## 5 Results

We trained each model on the MSE loss function using the complete dataset, i.e. all channels from all anatomical regions. Accuracy is measured by the average percentage of the error between the true and predicted images:

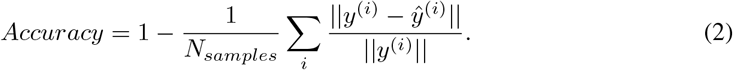

Table (1) lists the accuracy for each model and error metric. We see that the deep network outperforms the optimal linear decoder by roughly 15%. This metric improvement is also visible in the decoded images. A sample image comparing the presented image with the two model predictions is given in figure (4)

**Table 1:** Model Accuracy Computed using all *N*_*neurons*_ for linear, deep, and random (null) models. For the latter, pixel values of the predictions were randomly chosen to have same mean and unit variance as the target images from a Gaussian distribution.

| Model | Linear | Deep Network | Random Output |
| --- | --- | --- | --- |
| Train Accuracy | 40.0% | 55.5% | < 1% |
| Test Accuracy | 39.53% | 53.5% | < 1 a% |

**Figure 4:**
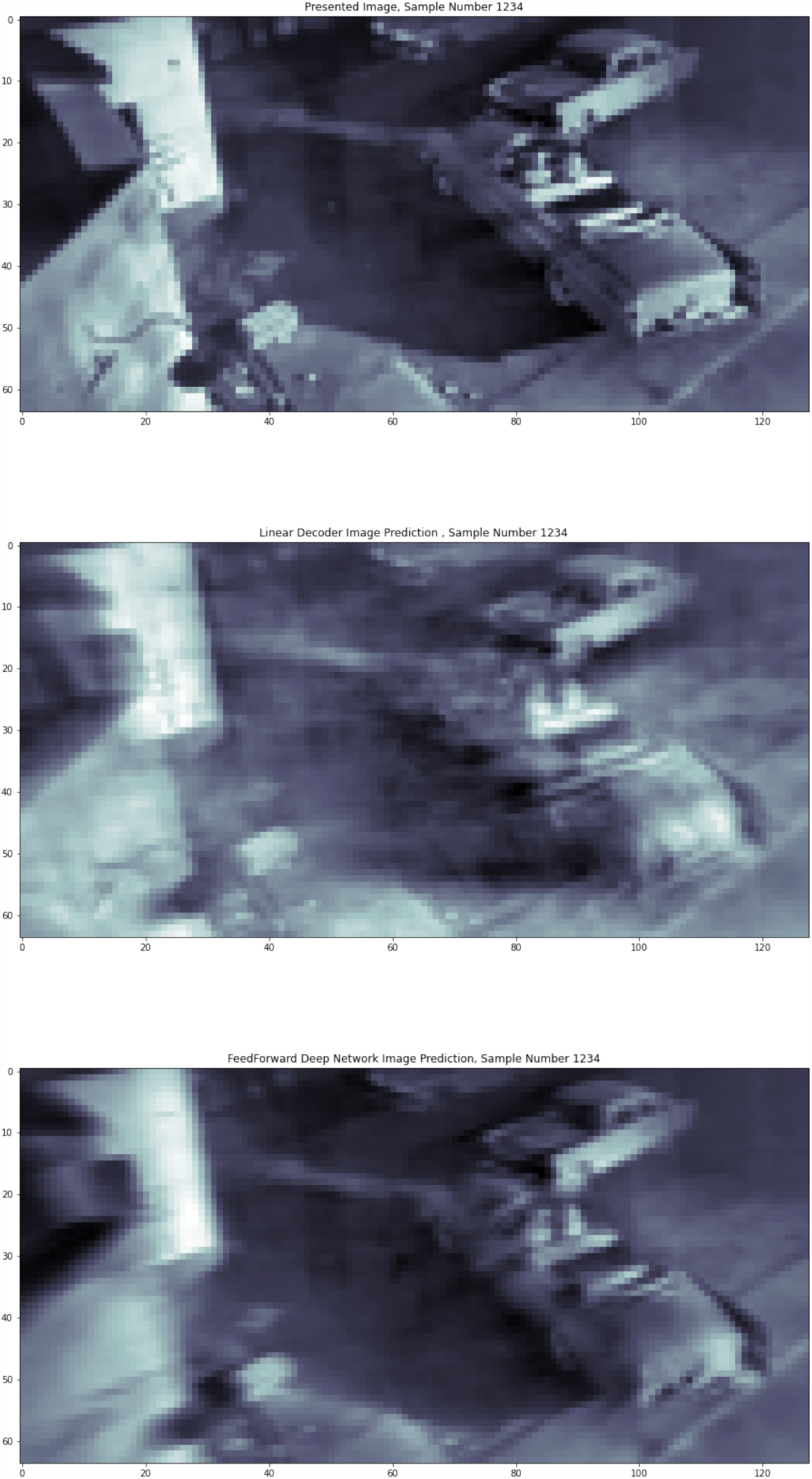
Comparison of Presented (Top) Image and Predictions Made by Linear (Middle) and Deep (Bottom) Decoders.

## 6 Conclusions & Future Work

We’ve shown that deep networks are more effective at decoding visual information from neural activity than their linear counterparts. This implies that a deep network will outperform a linear decoder when used in a visual brain-machine interface. Given the simplicity of the architecture and the expansive research in deep learning, future work could incorporate more sophisticated designs such as Convolutional Neural Networks(CNN’s), and various forms of Recursive Neural Networks(RNN’s) to process the sequential nature of temporal spike recordings.

## References

[1] Mikhail A Lebedev and Miguel AL Nicolelis. “Brain–machine interfaces: past, present and future”. In: TRENDS in Neurosciences 29.9 (2006), pp. 536–546.

[2] Daniel L Ruderman. “The statistics of natural images”. In: Network: computation in neural systems 5.4 (1994), pp. 517–548.

[3] William E Vinje and Jack L Gallant. “Sparse coding and decorrelation in primary visual cortex during natural vision”. In: Science 287.5456 (2000), pp. 1273–1276.

[4] Michael Weliky et al. “Coding of natural scenes in primary visual cortex”. In: Neuron 37.4 (2003), pp. 703–718.

[5] Emmanouil Froudarakis et al. “Population code in mouse V1 facilitates readout of natural scenes through increased sparseness”. In: Nature neuroscience 17.6 (2014), pp. 851–857.

[6] Brian Wandell and Stephen Thomas. “Foundations of vision”. In: Psyccritiques 42.7 (1997).

[7] Peter Dayan and Laurence F Abbott. Theoretical neuroscience: computational and mathematical modeling of neural systems. Computational Neuroscience Series, 2001.

[8] Torsten N Wiesel and David H Hubel. “Single-cell responses in striate cortex of kittens deprived of vision in one eye”. In: Journal of neurophysiology 26.6 (1963), pp. 1003–1017.

[9] Blake A Richards et al. “A deep learning framework for neuroscience”. In: Nature neuroscience 22.11 (2019), pp. 1761–1770.

[10] Daniel LK Yamins and James J DiCarlo. “Using goal-driven deep learning models to understand sensory cortex”. In: Nature neuroscience 19.3 (2016), pp. 356–365.

[11] David Sussillo et al. “Making brain–machine interfaces robust to future neural variability”. In: Nature communications 7.1 (2016), pp. 1–13.

[12] Francis R Willett et al. “High-performance brain-to-text communication via imagined hand-writing”. In: bioRxiv (2020).

[13] Eleanor Batty et al. “Multilayer recurrent network models of primate retinal ganglion cell responses”. In: (2016).

[14] Edwin M. Maynard, Craig T. Nordhausen, and Richard A. Normann. “The Utah Intracortical Electrode Array: A recording structure for potential brain-computer interfaces”. In: Electroencephalography and Clinical Neurophysiology 102.3 (1997), pp. 228–239. ISSN: 0013-4694. DOI: https://doi.org/10.1016/S0013-4694(96)95176-0. xURL: https://www.sciencedirect.com/science/article/pii/S0013469496951760.

[15] James J Jun et al. “Fully integrated silicon probes for high-density recording of neural activity”. In: Nature 551.7679 (2017), pp. 232–236.

[16] Joshua H. Siegle et al. “A survey of spiking activity reveals a functional hierarchy of mouse corticothalamic visual areas”. In: bioRxiv (2019). DOI: 10.1101/805010. eprint: https://www.biorxiv.org/content/early/2019/10/16/805010.full.pdf. URL: https://www.biorxiv.org/content/early/2019/10/16/805010.

[17] Visual Coding - Neuropixels. URL: https://portal.brain-map.org/explore/circuits/visual-coding-neuropixels.

[18] Marius Pachitariu et al. “Kilosort: realtime spike-sorting for extracellular electrophysiology with hundreds of channels”. In: BioRxiv (2016), p. 061481.

[19] Mark H Histed, Lauren A Carvalho, and John HR Maunsell. “Psychophysical measurement of contrast sensitivity in the behaving mouse”. In: Journal of neurophysiology 107.3 (2012), pp. 758–765.

[20] Weisi Lin and C-C Jay Kuo. “Perceptual visual quality metrics: A survey”. In: Journal of visual communication and image representation 22.4 (2011), pp. 297–312.

[21] Diederik P Kingma and Jimmy Ba. “Adam: A method for stochastic optimization”. In: arXiv preprint arXiv:1412.6980 (2014).

